# Resource or slot model in visual working memory: Are they different?

**DOI:** 10.1101/2024.02.08.579494

**Authors:** Fatemeh Hojjati, Ali Motahharynia, Armin Adibi, Iman Adibi, Mehdi Sanayei

## Abstract

When studying the working memory (WM), the ‘slot model’ and the ‘resource model’ are two main theories used to describe how information retention occurs. The slot model shows that WM capacity consists of a certain number of predefined slots available for information storage. This theory explains that there is a binary condition during information recall in which information is either wholly maintained within a slot or forgotten. The resource model gives a resolution-based approach defining a continuous resource able to be distributed among an unlimited number of items in the WM capacity. With newer hybrid models suggesting that WM may not strictly conform to one model, this study aimed to understand the relationship between the original models. By implementing correlational assessments of subjects’ performances in two different psychophysics tasks (analog recall paradigm with sequential bar presentation and delayed match-to-sample task (DMS) with checkerboard stimuli which are representative for resource and slot models, respectively), our study revealed significant correlations between WM performance (measured by DMS tasks) with recall error, precision, and sources of error (measured by sequential paradigm). Overall, the findings emphasize the importance of considering both models in understanding WM processes, shedding light on the debate between slot and resource models by demonstrating overlap in elements of both models.

## Introduction

Working Memory (WM) is a limited short term storage for temporary information retention and manipulation playing a critical role in multiple cognitive functions such as language comprehension, learning and reasoning [1, 2]. The WM capacity is a sensitive component influenced by different executive processes according to different neuropsychological models [3]. The conflict surrounding how this information is stored in WM has given rise to two popular theories: the ‘slot model’ and the ‘resource model’. The slot model conceptualizes WM capacity with a limited number of slots available in an all-or-none format for information storage. While, this model lacks a quality measure for resolution of recall, the resource model proposes a dynamic allocation of resources to memorized items, where memory precision decreases as the number of memorized items increases [4, 5].

The evaluation of WM typically involves various paradigms, such as delayed match-to-sample (DMS) tasks and analog recall tasks, each aiming to elucidate different characteristics and features of WM limits. While these tasks offer valuable insights, they exhibit distinct differences in their overall frameworks. For example, DMS tasks can interpret subject reactions based on correct or incorrect responses, assuming that either an item is fully maintained or forgotten without considering memory resolution. In contrast, analog recall tasks typically present a range of options for subjects to choose from, assuming internal and external noise influences memory recall. This raises the question of whether these tasks evaluate different aspects of the same concept or are they assessing distinct properties of WM.

While previously introduced WM paradigms were used to assess slot and resource models, recent computational models suggest that WM is not always confined within one of these traditional models, but rather has stimulus specific features and is not a solitary process [6]. This evidence suggest that strict categorization of visual WM between slot and resource models are less reflective of experimental data and a stimulus specific bias theory is more relevant [7]. Predicting performance outcomes using these tasks varies based of specific parameters. For instance, it relies on stimulus characteristics, object structure, complexity, and overall scene structure, all of which significantly impact WM performance [8-10]. With the goal to understand the correlation between WM precision and capacity, and the underlying similarities of the resource and slot model, we conducted this study. Subject performances in the DMS task with checkerboard stimuli and sequential paradigm with bar stimuli were correlated revealing intrinsic association between the two models.

## Methods

### Setting

Visual stimuli for task setup were generated with MATLAB software (MATLAB 2019a, The MathWorks, Inc, Natick, MA) and controlled by the Psychtoolbox 3 extension [11]. Subjects sat in a dimly lit room with a distance of ∼48cm from a cathode ray tube monitor (CRT, 15”, refresh rate of 75 Hz). A total of 30 healthy volunteers (7 females, 26.56 ± 4.61 years old, from 21 to 37 years old) were recruited for this study and enrolled in two visual WM tasks: analog recall paradigm with sequential bar presentation and a DMS with checkerboard stimuli.

### Sequential paradigm with bar stimuli

In the sequential task, each trial began with a central fixation point (0.26°) displayed for 2 seconds followed by presentation of a red, blue and green bar (pseudorandom order, 2.57° by 0.19°, Fig. 1A). The minimum angular difference between bars was 10 angular degrees and each bar was presented for 500ms and there was a 500ms delay (where a blank screen was displayed) between bars. Subjects were instructed to memorize the orientation of each bar. After presentation of the last bar, a vertical probe bar (in red, blue, or green) was presented to the subject. Participants were asked to adjust the orientation of the probe bar to one of the previously displayed bars with the same color (target bar) using a computer mouse. By clicking on the right button of the computer mouse, to confirm their decision, they received visual feedback showing the correct orientation of the target bar, their response, and the angular difference between their answer and the bar in question. We recorded the orientation of presented bars, subject’s response, and recall error (angular difference between target angle and subject response for each trial). Before beginning the main task, a 10-trial training block (with 1 bar, instead of 3) was used to familiarize the subjects with the procedure. We collected data from 6 blocks, each consisting of 30 trials (i.e., 180 trials per subject).

**Figure 1:**
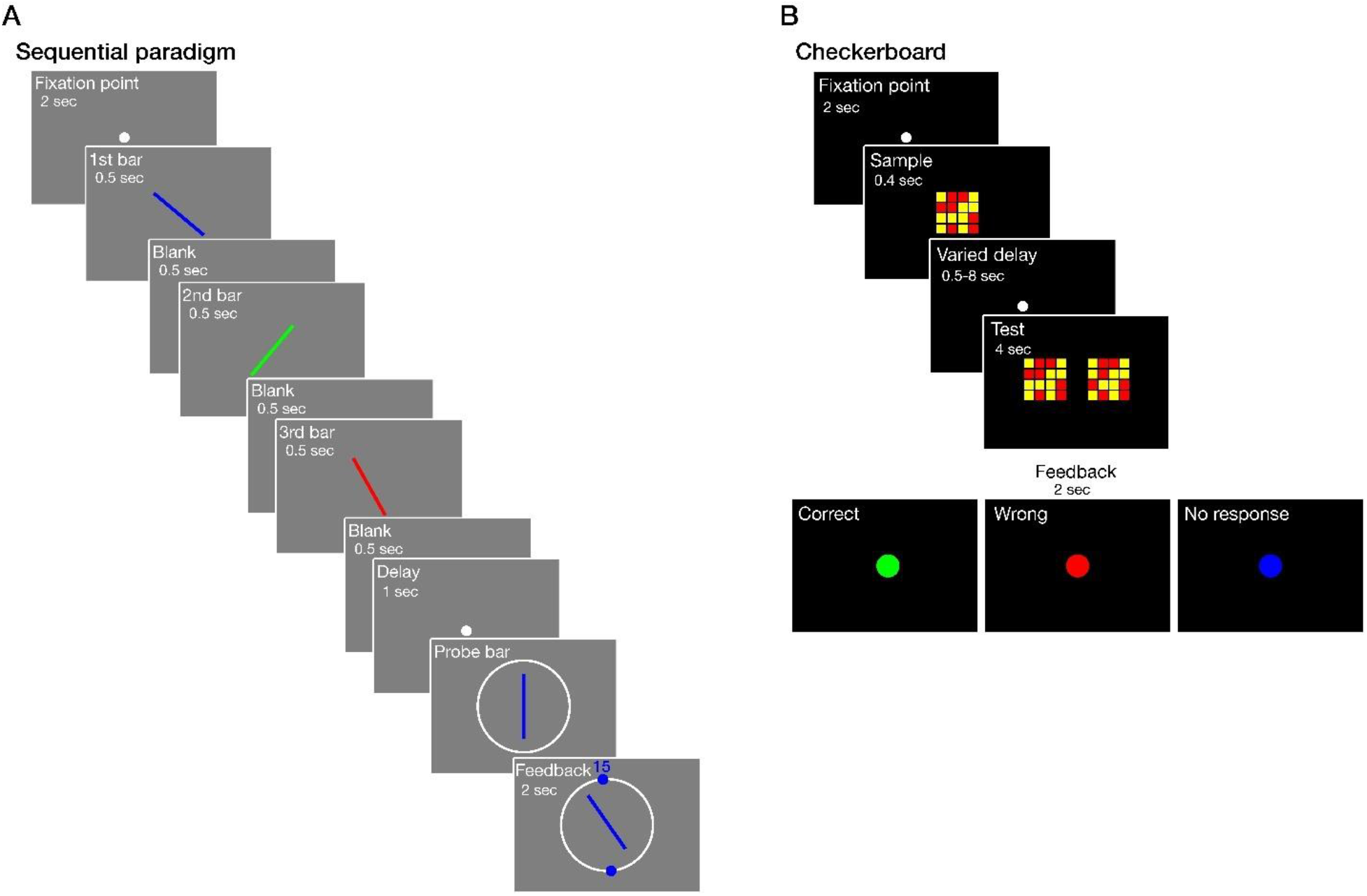
Schematic design of Working Memory (WM) tasks. A: In the analog recall paradigm with sequential bar presentation, subjects were asked to memorize bar orientations of three consecutively presented bars. After a 1s delay interval they were asked to match the probe bar to the angle of one of the previously presented bars with the same color. B: For the Delayed Match-to-Sample (DMS) task with checkerboard stimuli, subjects were asked to memorize a checkerboard pattern and after a random delay interval of 0.5, 1, 2, 4 or 8 seconds they were asked to select the correct pattern previously presented between two different checkerboard patterns.

### Delayed match-to-sample task (DMS)

For the DMS task, subjects were asked to look at a central fixation point (0.26°) for 2 seconds. After that a 4 by 4 checkerboard (4.83°×4.83°) was presented to the subjects (Fig. 1B). On each trial, 6 to 10 squares (out of 16) were yellow and the rest were red. Subjects were asked to memorize the pattern. After a delay of 0.5, 1, 2, 4, or 8 seconds, where only the fixation point was displayed, subjects were presented with two checkerboards. One was the same as the previously shown one (sample probe) and the other one had one yellow square swapped randomly with a red square. These two checkerboards were presented on the right and left side of the screen (3° from each other). Subjects were asked to report whether the right or the left checkerboard was the same as the sample stimulus by pressing the right or left arrow keys on the computer keyboard. Test checkerboards were kept on the screen until subject response or in a 4 second window. Visual feedbacks were given using a green disc (1.93°) for a correct, red for an incorrect response, and a blue one, if no response was given in 4 seconds. Like before, a 10-trial training block was used to familiarize the subjects with the procedure before beginning the main task. Each block consisted of 30 trials (5 repetitions for each delay) and each subject was enrolled in 6 trial blocks in total (i.e., 180 trials per subject).

### Statistical analysis

The statistical analyses were done with MATLAB 2023a. An analog report MATLAB toolbox from Bays lab was implemented calculating the circular mean for recall error for each subject [4]. Precision, defined as the reciprocal of circular standard deviation of recall error was calculated for each individual. To investigate the sources of error in recalling information, the Mixture Model was implemented. Using this model, the target proportion (Gaussian variability in reporting the orientation of the target bar), non-target proportion (swap error, Gaussian variability in misreporting the orientation of non-target bars) and, uniform proportion (random guessing) were calculated for each subject. Subject performance in the checkerboard task was defined as the ratio of correct responses to all given responses.

To calculate the correlation between tasks, we used Spearman’s correlation coefficient between performance in the DMS, and recall error and precision of the sequential task, separately. We also measured the correlation (Spearman’s) between the performance in DMS and the three sources of the error that we obtained from the Mixture Model. Furthermore, the correlation analyses between performance from each delay period (0.5, 1, 2, 4, and 8 seconds) with recall error, precision, and target, non-target, and uniform proportions of bar-specific responses were separately calculated.

Written informed consent was obtained from all participants. This study followed the latest update of the Declaration of Helsinki and was approved by the Iranian National Committee of Ethics in Biomedical Research (Approval ID: IR.MUI.MED.REC.1400.441).

## Results

A total of 30 healthy subjects (7 females, 26.56 ± 4.61 years old, from 21 to 37) participated in the analog recall task with sequential bar presentation and DMS task with checkerboard stimuli which is demonstrated in Fig. 1A and B. By conducting a Spearman’s correlation analysis, we found that subjects’ performance in the DMS task was negatively and significantly correlated with the mean recall error in the sequential task (r = −0.60, p < 0.001, Fig. 2A). We also found that performance in the DMS task was positively correlated with the precision of the sequential task (r = 0.60, p < 0.001, Fig. 2B). Furthermore, we found that while performance of the DMS task was positively correlated with target proportion (r = 0.59, p < 0.001, Fig. 2C), it was negatively correlated with non-target proportion (swap error, r = −0.55, p < 0.003, Fig. 2D). Performance did not show a significant correlation with uniform proportion (r = −0.22, p = 0.23, Fig. 2E).

**Figure 2:**
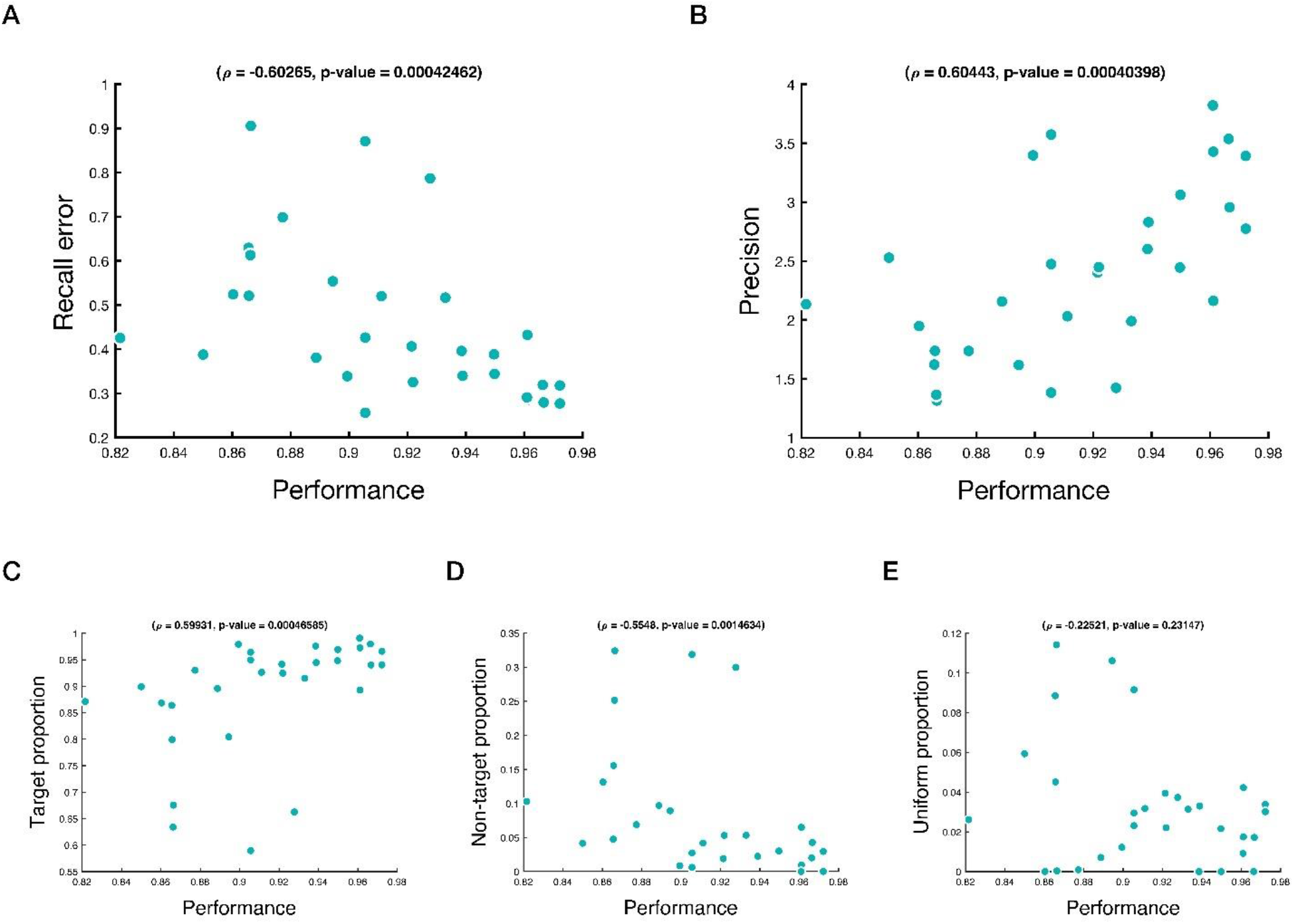
Correlations between parameters in analog recall paradigm with performance of DMS task. Correlation between (A) recall error, (B) precision, (C) target, (D) non-target (swap error), and (E) uniform proportions with DMS performance. Rho and p value of Pearson’s correlation are provided above each subplot.

A similar pattern was observed when we calculated the correlation between performance in DMS task for each delay, with recall error, precision, target proportion, non-target proportion, and uniform error for each bar order (Fig. 3A-E). We found that the performance in DMS task was more correlated with the recall error of the third presented bar (r = −0.59, p < 0.002), than the second (r = −0.56, p < 0.002), and first bar (r = −0.57, p < 0.001, Fig. 3A). The same pattern was observed for precision (3^rd^: r = 0.7, p < 0.001; 2^nd^: r = 0.46, p < 0.01; 1^st^: r = 0.45, p < 0.02, Fig 3B).

**Figure 3:**
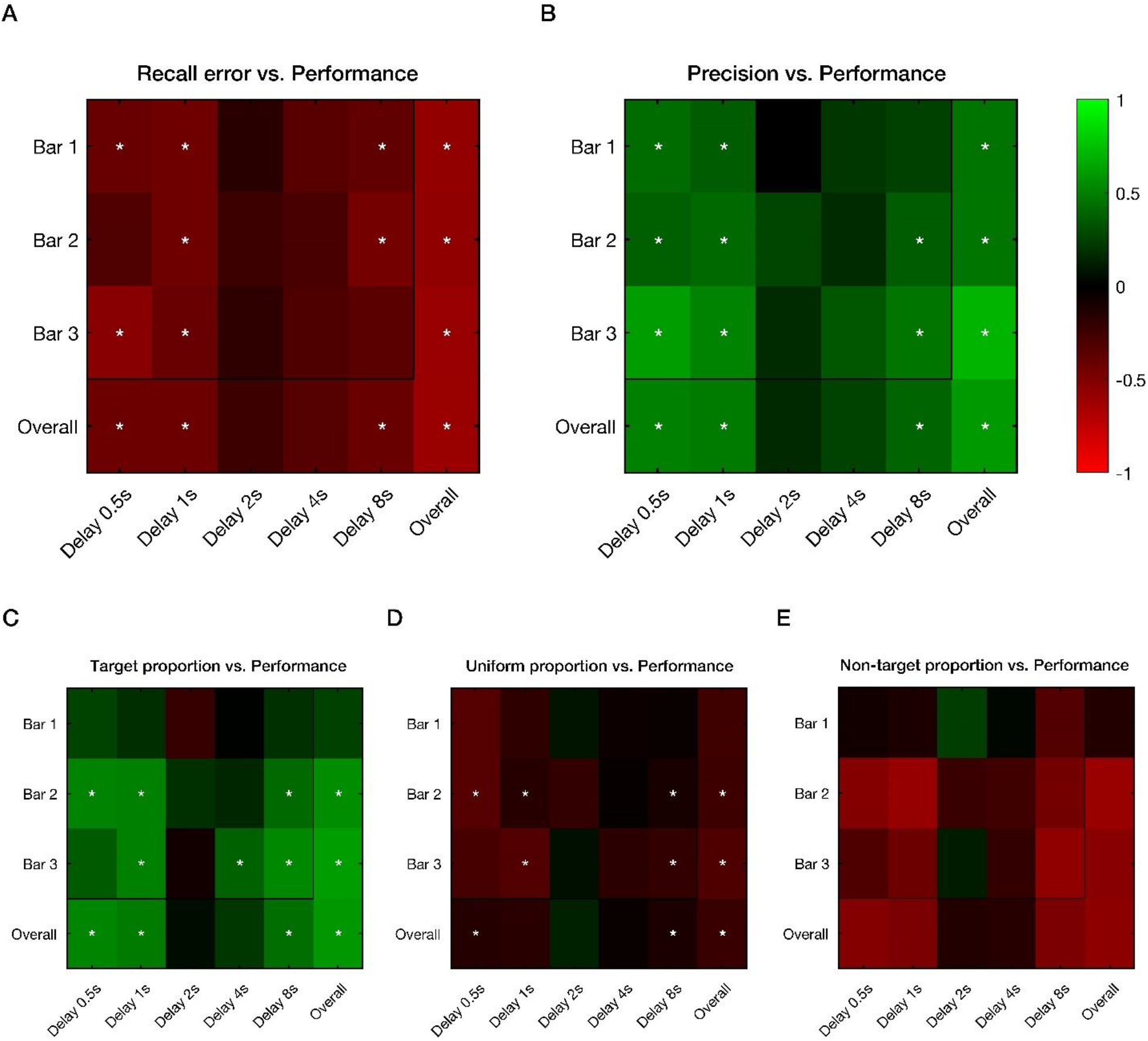
Correlation between bar orders in sequential paradigm vs. delay intervals in DMS. Heat map of Spearman’s correlation coefficient values between (A) recall error, (B) precision, (C) target, (D) non-target, and (E) uniform proportions, from bars 1 to 3 with checkerboard performance from five distinct delay periods (0.5, 1, 2, 4, and 8s). Asterisk shows significant correlation (p < 0.05), while green shades represent positive and red shades represent negative correlations.

## Discussion

The search for a comprehensive model explaining behavioral data from working memory (WM) tasks has led to the emergence of two prominent schools of thought: the slot-based model and the resource-based model. While each model possess unique properties capable of explaining various observation patterns in WM assessment, the discrepancies between them have been a subject of debate. Although recent studies have introduced hybrid theories, such as the categorical resource model, which incorporate features from both traditional models, correlative assessments have not been clearly implemented to absolve the differences of these theories [7]. Our correlational study aimed to unveil similarities and differences between these two mainstream models.

The moderate to high correlation observed between recall error and precision with DMS task performance, particularly noticeable in the third bar (the least challenging to memorize due to the shortest delay interval), highlights that when the memory load is lower in the sequential task, the results are more correlated to the less challenging task (i.e., DMS). This is complementary to a study by Zokaei et al., which compared digit span measures with a precision task and found significant correlations between performance in backward digit span (the more difficult condition in a slot-based model task) and precision in a resource model task [12]. In addition, our study had the benefit of showing the correlation pattern between these two tasks with higher temporal resolutions in DMS (from 0.5 to 8s) and sequential tasks (1 to 3 bars). These analyses showed no significant correlation in the 2s and 4s delay periods (Fig. 3, explained below), which shows the importance of considering full range of delay intervals.

The variation of correlation significance and power observed in our work is in line with a 2020 comparative analysis evaluating three visual WM tasks with distinct properties involving different types of stimuli and presentation formats to critique the comprehensive relevance of three prominent models (pure slot model, pure resource mode, and hybrid models) in explaining WM capacity [10]. While the slot-based model lacks the capacity to represent variability in memory resolution, continuous resource models assume internal and external noise according to signal detection theory. Although their findings were supportive of the pure resource model, regardless of task type, the joint model analysis showed performance in two tasks cannot be described with a single estimate of capacity or resource and it varies depending on simultaneous or sequential stimuli presentation. Resource distribution for information encoding and maintenance depends on information content and encoding conditions. They explain that studies supportive of the discrete slot model have overlooked base rate manipulation, set size variations (i.e., number of items asked to be memorized) and response bias (tendency to endorse a specific response). Therefore, the observed variability in correlation coefficients in specific delay intervals (2s and 4s) with recall error and precision could be described by different experimental settings in our study.

The absence of a significant correlation between uniform proportion (uniform error) and performance in the DMS task suggests that slot-based model tasks cannot be used to study uniform error. It is worth noting that the Mixture Model, introduced earlier by Bays et al., categorizes error patterns into three types: target, non-target, and uniform error. However, with the incorporation of the neural resource model (Stochastic sampling), the uniform error seems to be less relevant [13, 14].

In order to avoid any potential confound, our inclusion criteria were limited to individuals younger than 40 years old [15, 16]. However, this study had limitations which could be improved by conducting future EEG and functional MRI studies, along with behavioral paradigms, to distinguish between different models and pathways involved. Considering the impact of neuropsychological disorders such as Multiple Sclerosis, Alzheimer’s disease, and Parkinson’s on WM decline, which all require imaging modalities as a diagnostic step, integrating comparative analyses of imaging data with performance based tasks can help distinguishing different WM models in the future [17-19].

In conclusion, our study revealed a significant correlation between the resource and slot models, determining that the slot model is not necessarily outdated. This can serve as a confirmatory approach for when dealing with larger sample sizes and limited time, allowing reliance on classical models. However, when addressing sources of errors and their underlying features, classical slot models exhibit a weaker association with memory function.

## Data availability

Anonymized data will be available upon request from corresponding author. The corresponding author will consider the request against the data-sharing policy in the protocol and ethical approval of the study.

## Acknowledgment

This study was supported by Isfahan University of Medical Sciences (grant number: 2400104).

## Author contribution

Conceptualization: F.H, A.M, I.A, M.S. Paradigms development: A.M, M.S. Data acquisition: A.M, A.A. Statistical analysis: F.H, A.M, M.S. Data interpretation: F.H, A.M, I.A, M.S. Drafting original manuscript: F.H. Revising the manuscript: All authors. Funding acquisition: I.A. All the authors have read and approved the final version for publication.

## Code availability

The source codes for the analog report MATLAB toolbox are available at https://www.paulbays.com/toolbox/.

## Additional information

The authors declare no competing interests.

## Ethics

This study followed the latest update of the Declaration of Helsinki and was approved by the Iranian National Committee of Ethics in Biomedical Research (Approval ID: IR.MUI.MED.REC.1400.441).

